# A Multilevel Computational Characterization of Endophenotypes in Addiction

**DOI:** 10.1101/220905

**Authors:** Vincenzo G. Fiore, Dimitri Ognibene, Bryon Adinoff, Xiaosi Gu

## Abstract

Substance use disorders are characterized by a profound intersubject (phenotypic) variability in the expression of addictive symptomatology and propensity to relapse following treatment. However, laboratory investigations have primarily focused on common neural substrates in addiction, and have not yet been able to identify mechanisms that can account for the multifaceted phenotypic behaviors reported in literature. To investigate this knowledge gap theoretically, here we simulated phenotypic variations in addiction symptomology and responses to putative treatments, using both a neural model based on cortico-striatal circuit dynamics, and an algorithmic model of reinforcement learning. These simulations rely on the widely accepted assumption that both the ventral, model-based, goal-directed system and the dorsal, model-free, habitual system are vulnerable to extra-physiologic dopamine reinforcements triggered by drug consumption. We found that endophenotypic differences in the balance between the two circuit or control systems resulted in an inverted U-shape in optimal choice behavior. Specifically, greater unbalance led to a higher likelihood of developing addiction and more severe drug-taking behaviors. Furthermore, endophenotypes with opposite asymmetrical biases among cortico-striatal circuits expressed similar addiction behaviors, but responded differently to simulated treatments, suggesting personalized treatment development could rely on endophenotypic rather than phenotypic differentiations. We propose our simulated results, confirmed across neural and algorithmic levels of analysis, inform on a fundamental and, to date, neglected quantitative method to characterize clinical heterogeneity in addiction.

## Introduction

Substance use disorders are known to encompass a wide range of individual behavioral differences (i.e. phenotypes) in addiction development, maintenance and severity of symptoms, and treatment response (Everitt and Robbins, 2016). Previous investigations into the mechanisms underlying this heterogeneity of behaviors have identified two fundamental alterations that are correlated with a vulnerability in the development and severity of substance consumption (Garrison and Potenza, 2014; Jupp and Dalley, 2014; Belin et al., 2016). These neural and computational differentiations (i.e. endophenotypes) are respectively identified with a dysregulation of D2 receptors in the striatum (Morgan et al., 2002; Nader and Czoty, 2005; Dalley et al., 2007; Volkow et al., 2007; Flagel et al., 2010; Belcher et al., 2014; Flagel et al., 2014; Gould et al., 2014) and of learning rates (Gutkin et al., 2006; Piray et al., 2010). However, these endophenotypic differences are found across a wide spectrum of dissociable phenotypes, so that the same neural or computational mechanism is used to account for separable behavioral traits. For instance, striatal D2 dysregulation is found in individuals differing in terms of their impulsivity (Dalley et al., 2007; Volkow et al., 2007), social dominance (Morgan et al., 2002; Nader and Czoty, 2005; Gould et al., 2014), motor reactivity or preference for novelty (Flagel et al., 2010; Flagel et al., 2014) or sensitivity to rewards (Belcher et al., 2014). Each of these behavioral traits is separately correlated with development of addiction, but they do not necessarily coexist in the same individuals (cf. novelty seeking and impulsivity: Ersche et al., 2010; Molander et al., 2011; Belin and Deroche-Gamonet, 2012; Flagel et al., 2014). This mismatch between few known endophenotypic differences and a wide variety of multifaceted, dissociable, behavioral phenotypes suggests there are yet unknown neural and computational mechanisms that are responsible, alone or in interaction, for the reported behavioral differentiations. Finally, investigations into intersubject variability often emphasize the initial stage of addiction development (but see e.g.: Belin et al., 2008; Economidou et al., 2009; Pelloux et al., 2015). Yet, individual differences also emerge in response to treatments, resulting in diverse relapse patterns, among individuals showing similar severity of symptoms. These differences have not been so far addressed in neural or computational models.

To overcome these shortcomings, we propose a theoretical investigation into the interaction among ventral/dorsal cortico-striatal circuits and the associated behavioral control modalities. We have developed two models to simulate neural dynamics and algorithmic (or normative) choice selections in a multiple-choice task involving drug and non-drug rewards. Consistently with previous models, we assume addictive substances hijack the healthy reward prediction error signal (Schultz et al., 1997) by triggering extra-physiologic dopamine bursts (Nestler and Aghajanian, 1997; Koob and Volkow, 2016). These dopamine activities signal the presence of an aberrant unexpected reward, leading to the repetition of drug-related actions and escalation of consumption (Redish et al., 2008; Dayan, 2009). In our neural model, this process of reinforcement learning (RL, Sutton and Barto, 1998) is mediated by extra-physiologic changes in cortico-striatal connectivity weights (Hyman et al., 2006; Haber, 2008). These changes in turn aberrantly affect circuit gain and the stability of both ventral and dorsal cortico-striatal circuits, disrupting their respective role in encoding goal-directed behaviors (Balleine, 2005; Balleine and O’Doherty, 2010; Gruber and McDonald, 2012) and habitual responses (Yin et al., 2004; Balleine and O’Doherty, 2010). A similar effect is assumed for our algorithmic model, where over-evaluation of drugs and related RL affect the two control modalities, termed *model-based* and *model-free*, that loosely correspond to ventral/goal-oriented and dorsal/habitual implementations (Dolan and Dayan, 2013; Voon et al., 2017). As a result, and consistently with previous formulations of RL models of addiction (Redish et al., 2008; Piray et al., 2010; Gillan et al., 2016), both the planned evaluation of known action-outcome contingencies, represented in an *internal model* of the world, and the reactive immediate motor responses are biased towards drug-related selections.

Based on these assumptions, our models show that phenotypic differentiation in addiction development and treatment response can emerge as a function of a variable regulating the interaction between ventral and dorsal circuits or model-based and model-free control modalities. Our simulated results offer a proof of concept that this interaction is a candidate independent neural and computational mechanism underlying addiction vulnerability, putatively characterizing three different endophenotypes differing in the likelihood to develop addiction, severity of symptoms and treatment response. We suggest this neurocomputational mechanism could interact with previously described ones to generate the variety of dissociable behavioral traits reported in literature as associated with addiction vulnerabilities.

## Materials and Methods

In brief, we present two complementary models simulating endophenotypic differences and their effects on addiction development and treatment response. We tested our simulated agents under four conditions, and in environments granting free access to a substance of addiction, as usually implemented in laboratory studies. In particular, we compared our simulated phenotypic variability with the results described in a recent study investigating individual differences in rats self-administrating the stimulants cocaine or a designer drug, a dopamine- and mixed dopamine-norepinephrine reuptake inhibitor, respectively (Gannon et al., 2017). We selected this study because it highlights how different drugs, dosages and tasks result in different ranges of phenotypic differentiation. For instance, an initial acquisition phase, over a 10-day period, shows compulsive behavior developed in up to 75% rats self-administering cocaine and 87.5% of those exposed to the designer drug. Furthermore, under a condition of fixed ratio (=5) schedule, the study showed self-administration varied significantly among subjects. A subset of rat population, termed high responders, self-administered cocaine up to 60% more times in comparison with a different subset, termed low responders, depending on dosage (cf. figure 3 in: Gannon et al., 2017). Thus, our focus was on perturbing the balance between the dorsal/model-free and the ventral/model-based systems, to compare our simulated behavioral differentiations in drug consumption with the data reported in the chosen laboratory study.

The two models comprised a neural mass model that has been validated and described in the context of choice behavior and dopaminergic modulation (Baldassarre et al., 2013; Fiore et al., 2014; Fiore et al., 2016) and a normative or algorithmic model based upon standard RL schemes. In the neural model, addiction and treatment response were modeled through DA-dependent associative plasticity in both ventral and dorsal circuits. In the RL model, aberrant learning was modeled using a duplex of model-based and model-free schemes that competed for control over action selection. The model-based scheme entails learning a model of the environment (in the form of probability transition matrices among states) that was used to compute value functions under the Bellman optimality principle. The equivalent model-free scheme used prediction error based learning to directly acquire the value of state action pairs. Both neural and RL models were tested under four successive phases; 1) before exposure to drug (termed *pre-drug);* 2) learning of addictive behavior (termed *addiction*); 3) therapeutic interventions (termed *treatment)* that revert the learning of the previous phase. These are stylized interventions that simulate treatments prevalently affecting either goal-orented/model-based or habitual/model-free control systems. In the first case, the treatment is assumed to modify only the internal model of the environment and related selection of action-outcome contingencies performed in the ventral circuit. In the second case, the treatment represents a condition in which the model of the world of the agent remains mainly unaltered, but the acquired drug-related stimulus-response associations are disrupted, thus preventing the agent from exhibiting habitual responses (cf. Doll et al., 2009). Finally, 4) under the last condition we reinstated access to the simulated drug following each treatment (termed *relapse)*. The unique aspect of this complementary modeling approach is that neural and algorithmic models provide construct validation of each other, as process and implementation theories (i.e., synaptic and dynamical mechanisms) complement the normative principles formalized in the RL model.

### 1. Neural field model

#### Basic model architecture and parameterization

In cortico-striatal circuits, the signal processed in the cortex is conveyed towards its respective area of the striatum, processed in basal ganglia and finally relayed to the same cortical area where it originated, via thalamus (Haber, 2003; Draganski et al., 2008; Jahanshahi et al., 2015). Thus, despite diverging in terms of the information processed –e.g. sensorimotor or rewards and outcomes– these circuits are characterized by similar computational dynamics (Obeso et al., 2014). Temporal responses in recurrent neural networks co-occur with state transitions or input transformations that are often described in terms of energy landscapes (Fig. 1A-C). If multiple inputs or initial states generate transitions towards the same final state, this is termed *attractor state* (Amit, 1989). In recurrent networks such as cortico-striatal circuits, learning processes modulate the circuit gain, thereby affecting the strength of the attractor states and the overall stability of the system (Fiore et al., 2015).

**Figure 1.**
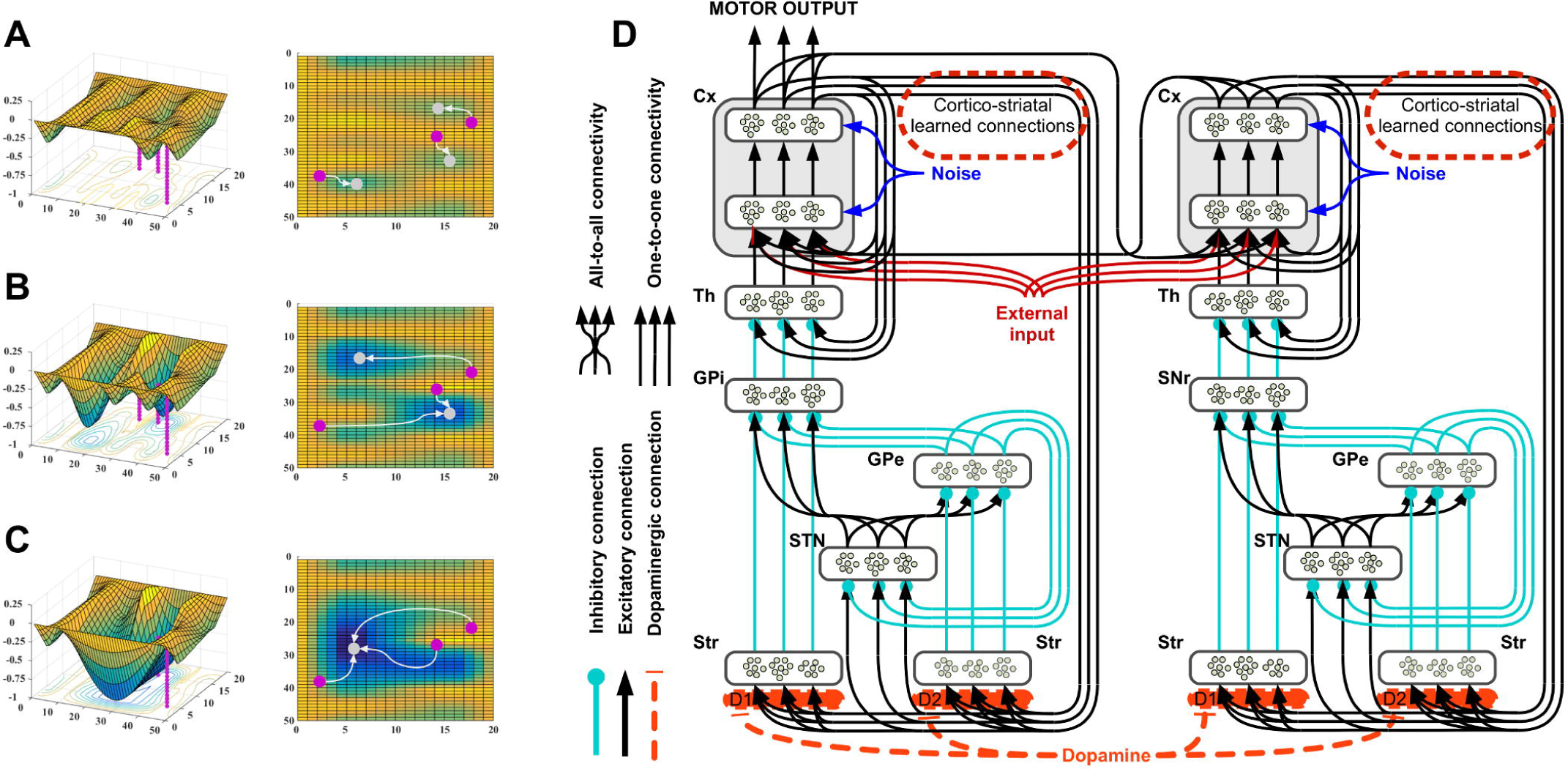
Illustrative representation of energy landscapes and neural architecture of the model. **A-C**. These representations of energy landscapes are meant to illustrate differences in the temporal responses provided by neural systems. Depending on the energy landscape, three arbitrary inputs (magenta circles) are transformed into different stable states (grey circles). Learning processes increase or decrease the strength of the connections among nodes in a network, thereby altering its energy landscape and reshaping temporal responses towards existing attractors. Attractors are defined as low energy states (bottom of the basins) at the end point of the temporal responses to multiple starting inputs. **A**. The landscape is characterized by multiple shallow attractors: these allow slow temporal responses, transforming multiple inputs into multiple weakly stable states. Noise and changes in the incoming input easily determine new responses towards different attractors. **B**. In this second illustrative configuration, steep and vast attractors characterize the energy landscape, allowing quick state transitions towards two equilibrium points. This new configuration is able to resist noise and minor changes in the incoming input, and, at the same time, allows a differentiation of inputs in two broad categories. **C**. Finally, the third energy landscape illustrates the presence of a parasitic attractor, exemplifying the condition of addiction: all inputs fall now at the bottom of a single steep basin. Under this condition, noise and changes in the incoming input determine temporal responses that keep falling in the same attractor, therefore preventing the system from executing different behaviors. **D**. Neural architecture used to simulate neural dynamics and behavior for the mean field neural model. The activity in the dorsal cortico-striatal circuit is responsible for the motor output of the system (left circuit), whilst activity in the ventral cortico-striatal circuit is responsible for goal selections (right circuit). The two systems bias each other via cortico-cortical connectivity and learning processes affect the weights of the connections between the two cortical outputs and the striatum in their corresponding circuits. The components in the architecture are labeled as follows: cortex (Cx), thalamus (Th), globus pallidus pars externa and interna (GPe and GPi), substantia nigra pars reticulata (SNr), sub-thalamic nucleus (STN) and striatum (Str), divided into two areas enriched by either D1 or D2 receptors.

We simulated the temporal responses in cortico-striatal circuits in a neural model developed in Matlab (illustrative representation of the neural architecture is represented in Fig. 1D). This neural model simulates mean-field activity (Deco et al., 2008) within multiple channels of both dorsal and ventral cortico-striatal loops. A continuous-time differential equation (equation 1) simulates changes over time (*τ_g_*) of the average action potential (*u_j_*) of a pool of neurons, and a positive transfer function (equation 2) converts this action potential in the final activation of the pool (*y_j_*). Finally, the plasticity of the connections (*w_ij_*) between cortex and striatum is characterized by DA-dependent Hebbian learning, corrected with a constant threshold (*th*) as defined in equation 3. The resulting rule strengthens the connections among all active nodes in the cortex and those active in the striatum, and weakens the connections among nodes showing opposite activation status.

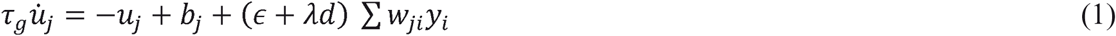

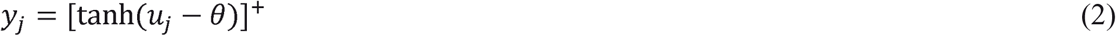

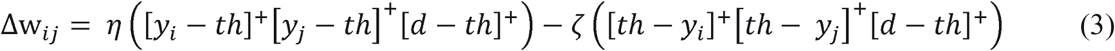

The input (∑ *w_ji_y_i_*), reaching each node in the neural network is modulated by two coefficients *λ* and *ε*. These regulate the ratio between the signal affected by the presence of dopamine release *d* and the amount of signal that is computed independent of dopamine release. For most units, the values of the two coefficients are set to λ = 0 and *ε* = 1, with the exception of the simulated striatal units, where these parameters are set to [ λ = 1.4, *λ* 0.2] and [λ = −0.5, ε = 0.6], to simulate the differential effect dopamine has, depending on the most prevalent receptor type (λ>1 and λ<0 for D1 and D2 receptors, respectively). Due to the different effects the dopamine receptors have on the activity of the simulated neurons, the drug-induced dopamine-dependent Hebbian learning significantly affects D1-enriched units in the striatum, whilst having negligible effects on D2-enriched units (Gerfen and Surmeier, 2011; Volkow and Morales, 2015).

#### Simulating different addiction phenotypes and treatment effects

Agents controlled by the neural model are immersed in a simplified environment and can select among three arbitrary actions or inactivity (cf. non stationary three armed bandit environment). The selection of the actions is carried out in the circuit simulating the dorsal cortico-striatal activity and it is considered completed if the neural activity of any of the units in the external layer of the simulated cortex (cf. Fig. 1) is maintained for at least 2 seconds. Ventral and dorsal circuits interact, both ways, via cortico-cortical connectivity. Therefore, the activity in the simulated ventral circuit biases action selection in the dorsal circuit and the selection of actions in the dorsal circuit biases the activity in the ventral circuit. To test our hypothesis about the effect these reciprocal biases have on choice behavior, we assumed cortico-cortical weights do not vary over time and we tested eleven combinations for the parameters determining their weights, as *w_ji_*=[0.02-0.2], [0.03-0.17], [0.05-0.15], [0.07-0.13], or [0.1-0.1] (and symmetrical). This spectrum of weights describes the strength of the biases between the two major circuits, thereby characterizing either a balanced condition or a dominance of one of the two circuits. We report the effects in terms of behavioral responses for these putative endophenotypes and test each of these with thirty noise seeds, random inputs and under four conditions, to allow within phenotype comparisons. The first condition, “pre-drug”, represents an assessment of behavior before any drug or reward is introduced, as the three available inputs randomly change their value to determine a non-stationary order of preferences. Under the second condition, termed “addiction”, one action is associated with the administration of a simulated addictive substance, triggering DA phasic responses and associated Hebbian learning in cortico-striatal connections of both ventral and dorsal circuits. For the third condition, termed “treatment”, we simulate the effects of deprivation coupled with one of two hypothetical treatments targeting either the dorsal or the ventral cortico-striatal circuits. The treatments are simulated reverting the learning process in either the dorsal or the ventral cortico-striatal circuit. The dorsal treatment brings back the pre-drug configuration in the dorsal circuit and keeps the configuration reached under the condition of addiction for the ventral circuit. The ventral treatment is achieved with the opposite intervention. Finally, under the fourth condition, termed “relapse”, we reintroduce access to the addictive substance, inducing relapse. Under this condition, relapse time is defined as the time required to reinstate the configuration of cortico-striatal weights found at the end of the *addiction* condition.

### 2. RL model

#### Basic model architecture and parameterization

In this model, we assume that the behavior of the agent relies on a hybrid model (Daw et al., 2011) that learns and computes the value of choices (actions, *a_t_*) under each condition (state, s_t_). Value is defined as a quantity that combines short and long term expected rewards and negative outcomes when a specific strategy of action is followed (policy, π). It is formally defined as:

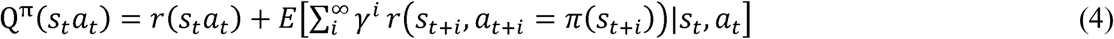

In equation (4), *r*(*s, a*) denotes the instantaneous reward received when action *a* is performed in state *s*. *γ* is a discount factor, comprised between 0 and 1, which defines the trade-off between immediate and long term rewards. The value of a state given the policy is defined as *V^π^*(*s*) = max_*a*_ *Q^π^*(*s,a*). For each environment there is an optimal policy *π*^*^(*s*), which maximizes the value *V^π^*^*^ (*s*) for every state (Sutton and Barto, 1998).

The environment can be completely characterized through the state transitions distributions *p*(*s*_*t*+1_ = *s|s_t_,a_t_*), and the expected rewards *E*(*r*|*s*,*a*) = *R*(*s*,*a*). These two functions together represent a model of the environment. Model-based behaviors compute Q^π^(*s_t_a_t_*) and the policy relying on such functions, at each state, following the Bellman equation (Daw and Dayan, 2014):

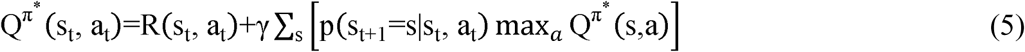

The model-based component learns the transition distributions and the expected rewards during the interaction with the environment. Thus, differently from other hybrid models (Daw et al., 2005; Keramati et al., 2011; Pezzulo et al., 2013), the quality of Q value estimation at any given moment depends on the experience the agent acquired up to that point in time. To compute value estimation (*Q_MB_*), this bounded (Gershman et al., 2015) component applies at each step the Bellman equation (5) a limited number of times (*N_PS_* = 50) to states sampled stochastically following a heuristic for efficient state update selection. The algorithm is an early-interrupted variation of the Prioritized Sweeping algorithm (Moore and Atkeson, 1993) with stochastic state update selection. Crucially, our model-based component does not accumulate the variations of Q values over time, and restarts the computation after each step (desJardins et al., 1999). This choice is meant to instate a plausible bounded rationality for our model which can account for the cognitive costs and ensuing limits of integrating old and new information about the environment, whilst updating and extending a complex plan to navigate it. This implementation was suitable for a bounded rational model-based component that shows controlled stochasticity of deliberation performances in non-trivial environments. This choice allowed to test the effects of the hypothesized endophenotypic differentiation in an environment characterized by higher degree of complexity in comparison with both the one chosen for the neural model and those described in the literature of RL models of addiction. In particular, we considered drug consumption to be associated with complex after-effects that make it difficult to predict the overall result of pursuing the related action course.

In comparison with other hybrid models such as Dyna and Dyna2 (Sutton, 1990; Silver et al., 2016), the proposed architecture does not share Q values between model-based and model-free components, nor it requires that the two processes share the same state representations. The two components separately represent their Q values and integrate them in a later phase. This decoupling is assumed to result in a more biologically plausible agent (Daw & Dayan 2014), and it is essential for the simulations of two separate treatments, essential requirement to establish a comparison with the behavior simulated with the neural model. In contrast with previous work using a hybrid Dyna-like architecture and Prioritized Sweeping algorithm, where the sharing of the Q-values explained the appearance of model based drug oriented behavior (Simon and Daw, 2012), in our model this model based addiction emerges in independent model-free and model based components. Thus, addiction behavior results from the joint effect of high reward (i.e. the drug), a limited number of stochastically selected policy updates and limited knowledge of the environment.

The model-free component has been implemented using the Q-Learning algorithm in tabular form (Watkins and Dayan, 1992). Q-learning updates initial state value estimations as follows:

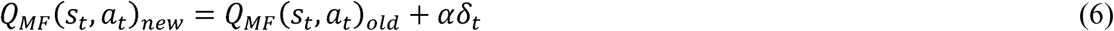

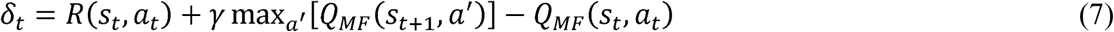

where *α* is a learning factor comprised between 0 and 1. Our hybrid model computes choice values in a fashion that balances model-free (MF in the equations) and model-based (MB in the equations) components depending on a parameter *β*. Six values (1, 0.8, 0.6, 0.4, 0.2, 0) are used for this parameter to simulate different endophenotypes, on a spectrum between purely model-based (*β*=1) and purely model-free (*β*=0) RL.

To allow exploration, the action to execute is selected randomly 10% of the times. This exploration factor is kept constant to support adaptation to a changing environment (Singh et al., 2000) and to simulate the continuous update of knowledge necessary to cope with ecological environments. The remaining 90% of the times, actions are determined by maximizing Q_MX_(s,a) in a strategy defined as ε-greedy (ε=.1). These values are produced by combining the values computed by the model-based and model-free components:

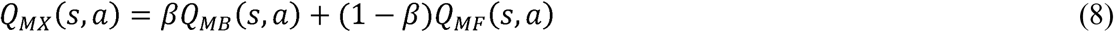

The choice for a fixed balance between model-based and model-free requires minimal assumptions on their interaction and has been used in recent reinforcement learning architectures (Silver et al., 2016).

#### Simulating different addiction phenotypes and treatment effects

In comparison with the simulations characterizing the neural model, for the RL model we could develop a more complex environment, to highlight how our endophenotypic differentiations could also affect the likelihood to develop addiction. This environment is characterized by a total of 20 states divided into four different types (Fig. 2): (i) healthy rewards (i.e. normal rewards that are not directly associated with drugs); (ii) neutral states (no reward or negative outcome); (iii) drug-related states, which give a high reward but are followed by multiple (iv) drug aftereffects, characterized by small negative outcomes. Similar to the neural model investigations, the agent deals with environment variations meant to simulate four phases of addiction: initial pre-drug phase (f1); addiction (i.e. the drug becomes accessible for the first time, f2); treatment (f3); relapse (i.e. second drug exposures, f4). Under the initial pre-drug phase (d_init_=50 steps), the agent does not receive any reward or negative outcome by entering the drug-related and aftereffects area, but a moderate reward is assigned (R_g_=1) by accessing the healthy reward state. Under the conditions of addiction and post-treatment addiction (d_tpy_=1000 steps), the agent can also receive a high reward, after accessing a drug-related state (Rd=10). The drug state always leads to a series of randomized state transitions among the aftereffects states (R_a_=-1.2) and simulates generic negative consequences associated with addiction. The agent can occasionally leave this aftereffect area of the environment (Fig. 2) to reach a neutral state, at the price of a further negative outcome (R_a_=-4). Under the treatment phase (d_tpy_=1000 steps), the drug-related state results in a negative outcome (Rdt=-1, see **Tables 1** and **2**, column f3), thus increasing the chances the agent stops pursuing this state. To allow for a comparison with the results in the neural model, we simulated a model-based and model-free treatment by manipulating the learning factor of the non-treated control modality, decreasing it: α_Ctpy_=0.01 * α. Under the condition of relapse, we measure the simulated time required by the agents to reach at least 95% of drug-related action preference as recorded under the addiction condition, after the drug is introduced again in the environment. This threshold is used to measure the percentage of agents relapsing, as well as the time required to complete the relapse, per endophenotype.

**Figure 2.**
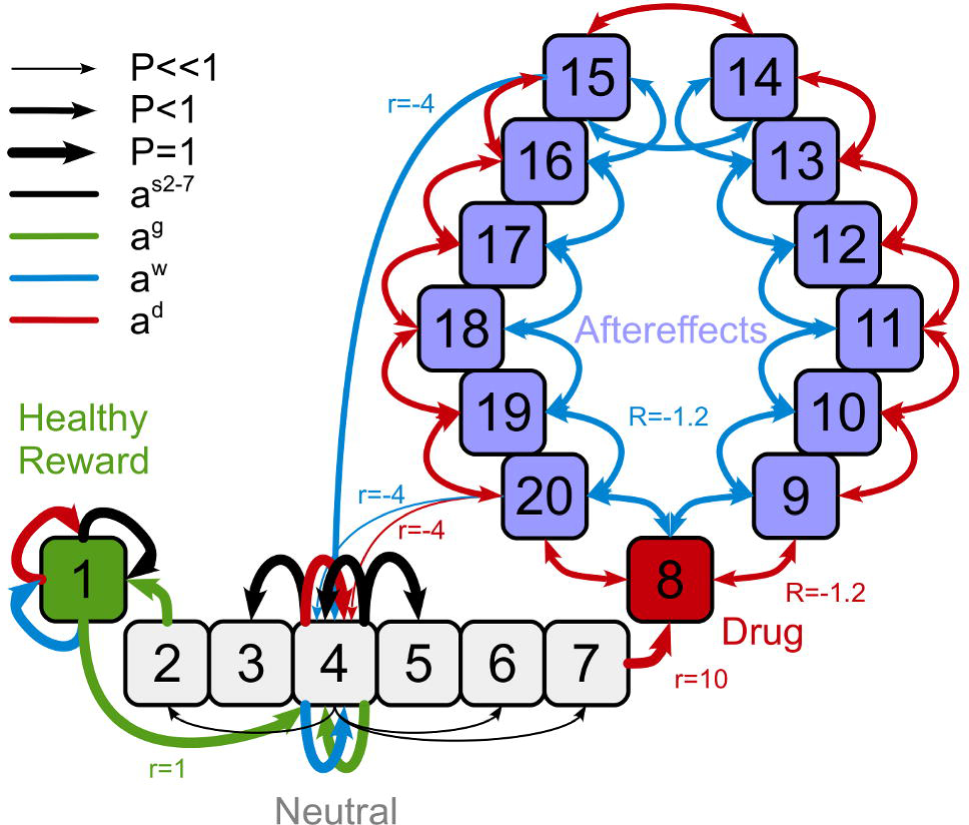
Illustrative representation of the environment used for the RL model of addiction. The states are disposed in a linear arrangement: on one extreme is a healthy reward state (1), on the opposite side a drug state (8) followed by fourteen aftereffects states (9-22). Healthy reward and drug states are separated by 6 neutral states (2-7). The agent can traverse between nearby neutral states. From the two borders of the central segment of neutral states, an agent can enter the healthy reward state (from state 2), securing a moderate reward (Rg=1), or the drug state (from state 7), receiving an initial high reward (Rd=10, under the condition of addiction) and a series of sparse but temporally extended negative outcomes, characterizing the aftereffects states. The presence of negative outcomes makes entering the drug and aftereffects area suboptimal under all experimental conditions (see optimal policy in **table 3**). From both the goal state and the drug/aftereffects segment the agent is then returned to the middle of the neutral segment. In this representation, we explicitly portray the transitions related to states 1 (healthy reward), 4 (neutral), and 15 and 20 (drug aftereffects) for illustrative purposes. Line width represents related transition probability value. Line and text color represent the action class (a_s_, a_g_, a_w_, a_d_). Neutral states are navigable with actions a_s2-7_ which are deterministic for adjacent state while have high chance of failing for distant states. From the neutral states the agent can reach: (i) the healthy reward, if executing action a_g_ when in state 2; and (ii) the drug state (8) and aftereffects area (state 9 to 22), if executing action a_d_, when in state 7. From the healthy reward area the agent can issue again a_g_, receiving a reward of 1 and going back to the center of the neutral area, state 4. By entering the drug area, the agent receives a reward of 10. Action results in the drug/aftereffect area are probabilistic: the agent can reach a nearby state in the area or leave the area and reach the center of the neutral state. Leaving the drug/aftereffects area has a cost of -4, whereas every other transition inside the area costs -1.2. For a full description of transitions and their probability distribution in the environment (see **Tables 1-2,4-5**).

## Results

### Simulations from the neural field model

Under all conditions, the three stimuli randomly change every few seconds, putatively representing a dynamic fluctuation of values associated with perceived cues in a non-static environment. Optimal behavior requires the agents to rapidly adapt to these changes, transiently triggering the motor response associated with the most valuable cue (cf. Fiore et al., 2016; Hauser et al., 2016). Under pre-drug condition, dorsal and ventral circuits perform unbiased selections, collaborating in the generation of a near-optimal sequence of motor selections. All eleven endophenotypes show uniform distributions of action selections, complying with the random distribution of the inputs configurations (Fig. 3A). This control condition allows the simulated network to generate transient temporal responses that couple multiple initial states with multiple stable states, in a transient winner-take-all or *winner-less* competition (Rabinovich et al., 2006; Afraimovich et al., 2008).

**Figure 3.**
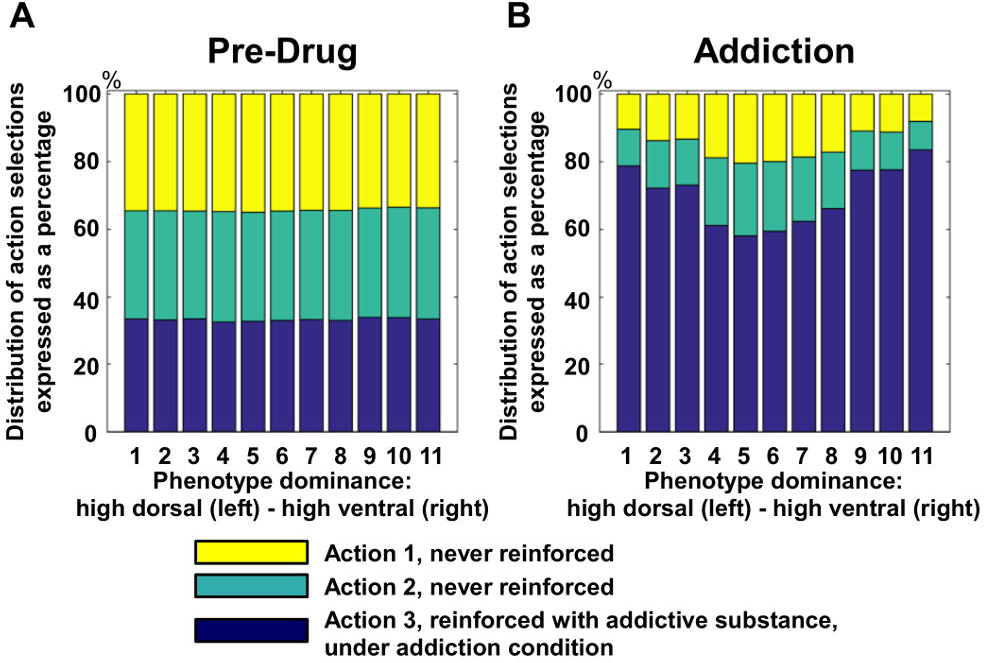
Histograms show how the distribution of simulated action selections changes depending on the endophenotype (11 structural variations in cortico-cortical connectivity). 30 random seeds/inputs are used per phenotype, tested under two conditions: pre-drug (A) and addiction (B). The three colors represent the occurrence of selections of three arbitrary actions. Under the pre-drug condition, no reward is provided and action selections are triggered by random fluctuation in values of competing sensory inputs. The simulations show the agents adapt to the changes in sensory stimuli and therefore exhibit a near-uniform distribution of action selections. Conversely, under the condition of addiction, the action represented in blue is associated with administration of the simulated drug, triggering DA-dependent Hebbian learning and consequently over-selection. Under addiction, the differences among endophenotypes clearly emerge in the selection frequency of the action leading to drug consumption. Asymmetric control (endophenotypes 1-3 and 9-11) leads to a stronger over-selection in comparison with balanced control (endophenotypes 4-7), despite identical learning processes and reward encoding.

Under the condition of simulated addiction, one of the actions is associated with drug administration (Fig. 3B, values represented in blue). Substance use triggers phasic dopamine bursts, leading to Hebbian learning in cortico-striatal connections of both dorsal and ventral circuits (equation 3). In recurrent networks, circuit gain increases as a direct function of the weights of reentrant synapses (Amit, 1989). A dopamine response triggered by healthy unexpected rewards would create a bias towards the selection of the reinforced motor response to a perceived cue (Cohen and Frank, 2009; Grahn et al., 2009). However, drug consumption triggers extra-physiologic dopamine-dependent learning, resulting in aberrantly high circuit gain, and compromising the ability of all affected circuits to discriminate among different inputs and produce temporal transitions towards multiple stable states. The cortico-striatal circuits become over-stable and resistant to perturbation caused by a change of input or by noise (Fiore et al., 2014) as they are dominated by *parasitic attractors* (Hoffman and McGlashan, 2001) (Fig 1C). In the ventral cortico-striatal circuit, a parasitic attractor sets and maintains the selection of drug-related goals or outcomes, biasing the action-outcome assessments required for planning. In the dorsal circuit, the same process determines over-stable selections of the reinforced motor behavior, generating reactive responses and habits. Importantly, the learning process simulated in our neural model led to the generation of parasitic attractors in both circuits across all endophenotypes, as all agents eventually reached a fixed threshold in cortico-striatal neural plasticity. Despite the generation of a form of compulsive drug seeking behavior across all endophenotypes, we observed significant differences in motor response patterns as a function of the balance between ventral and dorsal circuits. Specifically, the endophenotypes characterized by unbalanced dorsal or ventral control (i.e. Fig. 3B, endophenotypes 1-3 and 9-11) expressed distributions of motor selections that were significantly more compromised by drug-related aberrant rewards, in comparison with balanced endophenotypes (i.e. Fig. 3B, endophenotypes 5-7). The presence of identical learning processes, and the associated attractor formation in both ventral and dorsal circuits, ascribes all phenotypic differences univocally to the only remaining independent variable, which controls cortico-cortical connectivity and therefore the strength of the biases between circuits. Unbalanced agents were characterized by more frequent drug-related selections as actions leading to drug consumption were selected more frequently than in balanced endophenotypes, in a range between +3% and +45%. This result would identify all phenotypes within the limits of individual differentiation described in the study chosen for behavioral comparison (Gannon et al., 2017).

Next, we investigated how the simulated endophenotypes behaved under the conditions of treatment and relapse. We measured the frequency of drug-related action selections under the conditions of addiction and treatment (Fig. 4A-B). Both ventral (goal-oriented) and dorsal (habitual) treatments effectively reduced the number of actions associated with drug consumption, in comparison with baseline addiction. However, the dorsal treatment was more effective for dorsal-dominated endophenotypes and the ventral treatment was more effective for ventral-dominated endophenotypes. These endophenotype-specific treatment effects were further confirmed by our analysis of individual differences under the relapse condition (Fig. 4C-D): dorsal treatments were more effective in elongating time to relapse for dorsal-dominated endophenotypes, whereas ventral treatments were more successful in delaying relapse for ventral-dominated endophenotypes. Together, both measures showed that our simulated treatments have differential effects depending on the balance between dorsal and ventral circuits. Importantly, these differences emerged only after the treatment was applied, where a pretreatment comparison between compulsive behaviors expressed by the opposite unbalanced endophenotypes (i.e. ventral-dominant or dorsal-dominant) did not show any significant difference in choice selections (cf. Fig. 3B, endophenotypes 1-3 and 9-11).

**Figure 4.**
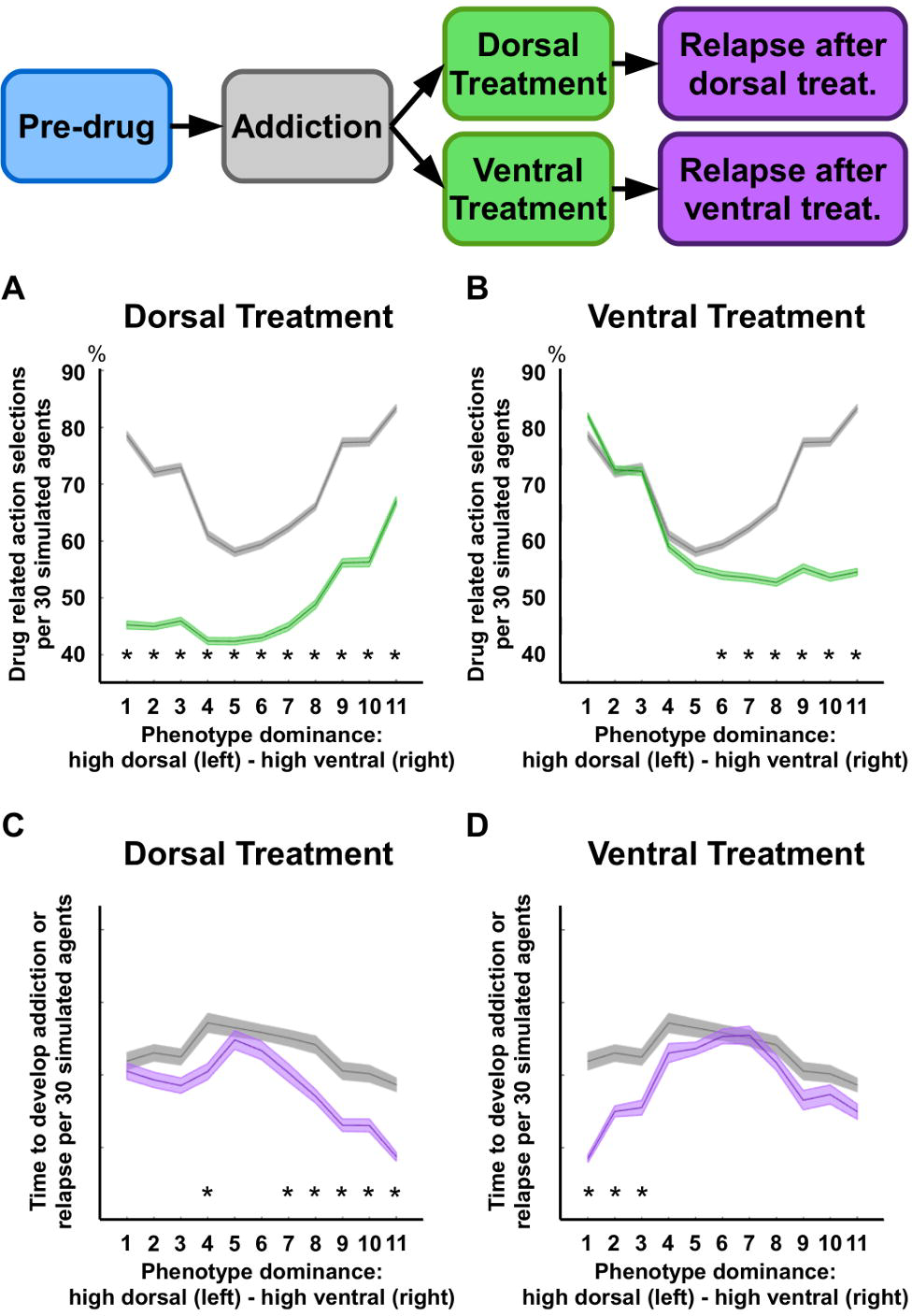
Shaded error bars report mean and standard error for 30 simulated agents across 11 endophenotypes (structural variations in cortico-cortical connectivity). Panels A and B (top panels) show the selections of actions leading to substance consumption, as a percentage of the overall number of action selections. In the first case (A) we compare the values recorded under the condition of addiction with those recorded under the condition of dorsal treatment jointly with abstinence (i.e. drug-related actions do not trigger self-administration of a drug and the treatment targets the dorsal circuit). In the second case (B) the comparison involves addiction and ventral treatment (treatment targeting the ventral circuit, during abstinence). Panels C and D (bottom panels) compare the simulated time required by the 11 endophenotypes to reach an arbitrary threshold of cortico-striatal connectivity under the condition of addiction and under the condition of relapse after either dorsal (C) or ventral (D) treatment. Within the time of a simulation run, all simulated agents reached the addiction threshold under all conditions here reported. The two treatments are simulated by restoring either the dorsal/motor (A-C) or the ventral/outcome circuit (B-D) to the configuration characterizing the pre-drug condition. The percentage of the action selections shows the dorsal treatment is more effective in endophenotypes characterized by high dorsal dominance (A), whereas the ventral treatment only has an effect in endophenotypes characterized by high ventral dominance (B). Similarly, dorsal and ventral treatments result in long relapse times in endophenotypes characterized by high dorsal and high ventral dominance, respectively. (*) indicates significant difference: p<0.05.

### Simulations from the RL model

By simulating explicit negative outcomes associated with drug consumption, the RL model allowed to measure the likelihood each agent has to develop addiction, as a function of its endophenotype. In our analysis, addiction is defined as a behavior leading to drug selections more frequently than the healthy alternative reward, under the addiction phase. The mean percentage of these *addicted* agents (over 300 runs) was 43.05%, across endophenotypes, which is consistent with the percentage of rats developing compulsive self-administration of cocaine, as reported in the reference study (~40% over a period of 5 days, cf. Gannon et al., 2017). Importantly, when considering endophenotype differentiation, the percentage varies significantly: 60.3% for *β*=0, 40.3% for *β* =0.2, 30.1% for *β* =0.4, 36.7% for *β* =0.6, 39.3% for *β* =0.8, and 51.6% for β=1 (Fig. 5A-B). This phenotypic differentiation is consistent with well-established data from animal models. For instance, rat strains selectively bred for either high or low voluntary running differ in the likelihood to develop addiction when given free access to cocaine (respectively ~35% and ~60% of each strain develop addiction over a period of 5 days, cf. Smethells et al., 2016). Free access to substances of abuse does not necessarily lead to compulsive behaviors (Piazza et al., 1989; Belin et al., 2011), as addiction varies as a function factors such as exposure extent, amount of drug delivered, and associated negative effects (Pelloux et al., 2007; Jonkman et al., 2012). Our simulations suggest that endophenotypes with lower chances of addiction are characterized by balanced control modalities. Note that an optimal agent, knowing the environment structure and being able to compute the long-term effects of drug, will never select drug-states (**Table 3**).

**Figure 5.**
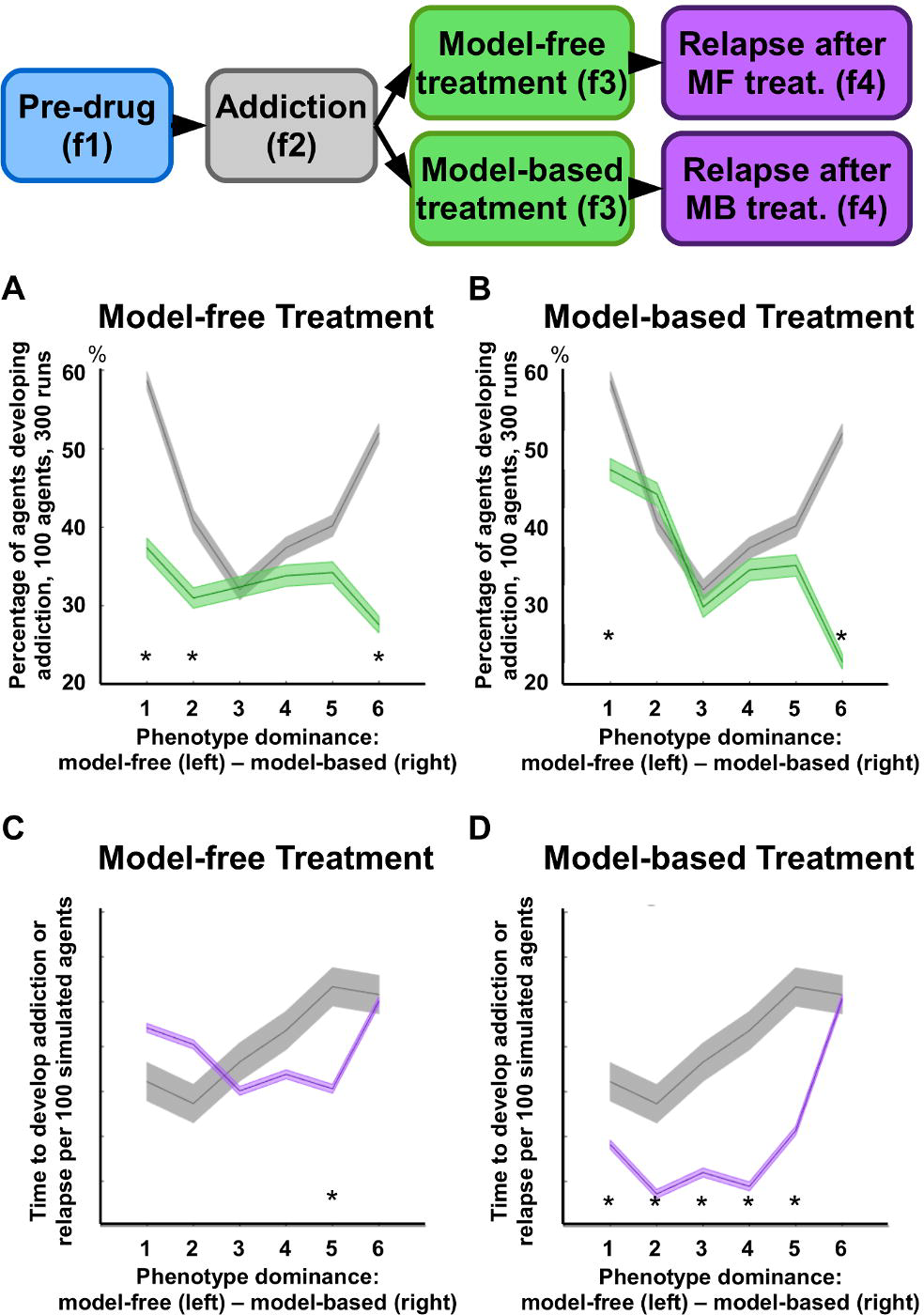
Shaded error bars report me an and standard error for _ 100 simulated agents across 6 endophenotypes (differential balance between model-based and model-free control modalities, P=[0, 0.2, 0.4, 0.8, 1]). Panels A and B show the percentage of agents developing addiction (i.e. drug-related choices are more frequent than healthy reward-related choices), per endophenotype, under the condition of addiction and treatment. In the first case (A) the comparison involves data recorded under the condition of addiction and those recorded under the condition of model-free treatment. In the second case (B) the comparison involves the condition of addiction and model-based treatment. Panels C and D illustrate the simulated time required by the 6 endophenotypes to reach 95% of action preference towards the drug state, in comparison with action preference recorded under the condition of addiction (f2). In the first case (C) the comparison involves the conditions of addiction and relapse after model-free treatment, whereas in the second case (D) the comparison involves the conditions of addiction and relapse after model-based treatment. In terms of action selection ratio, the simulated results show both treatments have a significant effect only on those phenotypes characterized by strong unbalance of control (A-B). In terms of relapse, the results show the model-free treatment is on average more successful than the model-based one, as 5 endophenotypes show no significant difference between the condition of addiction and post-treatment addiction (i.e. the time required to relapse is not significantly different than the time required to develop addiction the first time). Each endophenotype, or parameter selection, was simulated 100 times across the four phases (3050 steps per simulation). Results depend on the statics of the environment, but over similar environments the results were qualitatively similar. (*) indicates significant difference: p<0.05.

Finally, the simulations suggest that the hypothetical treatment targeting model-free control is the most effective, reducing the likelihood to pursue drug-related behaviors for all endophenotypes (Fig. 5A). In contrast, the model-based treatment appears to be less effective for all endophenotypes, with the exception of the purely model-based one (*β*=1) (Fig. 5B). Under the condition of relapse, our data confirm that the simulated treatments significantly differ in their effectiveness across the proposed endophenotypes, also suggesting the treatment targeting model-free control is the most successful in prolonging relapse time (Fig. 5C-D). Relapse time after model-free treatment is mostly similar to the time required to develop addiction behavior before any treatment (Fig. 5C). At the opposite side of the control spectrum, the model-based treatment shows a positive effect only for the purely model-based endophenotype. All remaining endophenotypes show relapse times significantly shorter than those recorded the first time addiction is developed (*β*=1; Fig. 5D).

## Discussion

Individual differences in stress and anxiety responses (Dilleen et al., 2012; Jimenez and Grant, 2017), social dominance (Morgan et al., 2002; Covington and Miczek, 2005), aggressive temperament (McClintick and Grant, 2016), preference for saccharine (Carroll et al., 2002), sensation or novelty seeking (Suto et al., 2001; Nadal et al., 2002; Belin et al., 2011; Flagel et al., 2014), impulsivity (Perry and Carroll, 2008; Verdejo-Garcia et al., 2008; Dalley et al., 2011), and sensitivity to rewards (Belcher et al., 2014) have all been found in both animal models and clinical studies in humans to be associated with addiction vulnerabilities, and in particular with the likelihood to develop and maintain addiction, or to resist to treatment (Piazza et al., 1989; Belin et al., 2016; Everitt and Robbins, 2016). However, investigations into the mechanisms underlying this phenotypic differentiation in addiction has so far revealed few neural or computational candidates, which are found associated with diverse and dissociable behavioral traits. An important example is represented by the endophenotypic differentiation reported in the expression and reactivity of striatal D2 dopaminergic receptors, which is found across the traits of impulsivity (Dalley et al., 2007), social dominance (Morgan et al., 2002), novelty preference (Flagel et al., 2014) and sensitivity to rewards (Belcher et al., 2014). The overlap of this endophenotypic trait across multiple, non-coexisting, phenotypes associated with addiction vulnerabilities suggests other neural or computational mechanisms have yet to be identified to allow accounting for the reported variety in behavioural traits.

Here we have presented a neural field model, augmented by an RL model, to expand on existing neuropsychological and computational accounts of addiction. Our models propose a theoretical investigation into the interaction among cortico-striatal circuits or behavioral control modalities, and the effects this interaction has on addiction development, and treatment response. As classically described in previous models (Redish, 2004; Redish et al., 2008; Dayan, 2009), we have assumed that over-evaluation of a drug leads to aberrant dopamine release and associated over-learning in multiple DA targets (Bjorklund and Dunnett, 2007). In the neural field model, this mechanism results in the dysregulation of the circuit gain and associated dynamics of both ventral and dorsal cortico-striatal circuits. In the integrated model-based and model-free RL model, sequential choice behavior is confounded by the presence of a high immediate reward (drug state), which leads to misrepresent the negative outcomes following drug consumption, if their distribution across states and time is sufficiently complex to escape the capabilities of the agent to correctly represent the environment (Doll and Daw, 2016; Sadacca et al., 2016). We found that both models jointly indicate that the balance between neural circuits or behavioral control modalities is a candidate neurocomputational mechanism characterizing endophenotypes in addiction. The neural and RL models converge in suggesting that individuals characterized by balanced behavioral control between reward-seeking or planning (ventral circuit/model-based) and reactive or habitual responses (dorsal circuit/model-free) would have a reduced chance to develop addiction and decreased severity of symptoms if developing addiction. We propose this neurocomputational mechanism may be interacting with other known endophenotypic differentiations, such as alterations of D2 receptors in the striatum (Morgan et al., 2002; Nader and Czoty, 2005; Dalley et al., 2007; Volkow et al., 2007; Belcher et al., 2014; Flagel et al., 2014) or differences in learning rates (Gutkin et al., 2006; Piray et al., 2010), to generate the multifaceted behavioral traits that have been reported in literature to be associated with addiction vulnerabilities.

In our neural model, ventral and dorsal circuits are mostly in phase in their selections under the pre-drug condition, exhibiting synchronous transient stability of neural activity and enhancing the overall ability of the system to adapt to changing stimuli (i.e. the two circuits adapt to the input changes with a similar pace and synchronize in their selection). Under the condition of addiction, the two circuits are mostly pulled towards the parasitic attractor state associated with drug consumption, and they occasionally select the competing non-drug stimuli. If only one of the two systems performs a selection outside of the attractor, the difference in selection generates a dissonance or interference. In neural endophenotypes characterized by unbalanced control, this dissonance is solved by one circuit taking the lead, so that both systems eventually converge on the selection of the dominant circuit. These dynamics result in limited opportunities to generate non-drug responses to the external stimuli, as they can only be generated by the dominant circuit. Conversely, in balanced control endophenotypes, if any of the two circuits ignores the drug-stimulus and selects a competing option, the resulting dissonance can trigger a state transition pulling out the parasitic attractor states associated with substance use. The endophenotypes in our simulations vary only in the parameters regulating the balance between circuits, as dopamine-driven learning processes established between cortex and striatum (equation 3) do not vary across endophenotypes, resulting in identical habit formation and drug-related biases in the outcome representations. Thus, our proposed phenotypic differentiation does not interfere with the usual role ascribed to the ventral and dorsal circuits as respectively implicated in the initial reward-seeking phase in addiction (Belin and Everitt, 2008; Willuhn et al., 2012) and the subsequent consolidation of stimulus-response, habitual, association (Everitt and Robbins, 2013, 2016). However, our simulated dynamics show that, after addiction is developed, systemic over-stability can be reduced or further enhanced, depending on the cortico-cortical biases between cortico-striatal circuits. In turn, this modulation of system stability can foster or further impair input discrimination and motor response versatility, affecting addiction symptomatology. As a result, our neural model shows phenotypic variability emerging after the presentation of the reward simulating the drug and addiction is developed, in a gradient of over-selection of drug-related actions.

In the RL model, we investigated whether the balance between model-based and model-free modalities would also increase the robustness of the system against the selection of drug-states in a more complex environment and in presence of explicit negative outcomes. Similar to the neural model, a system with balanced control modalities introduces more diversity in action selection during exploration, reducing (yet not cancelling) the chances of developing maladaptive reactive responses. This increased diversity and overall reliability are likely to be induced by a higher redundancy and diversification of the system. While both components may fail, the causes of failures are not necessarily correlated. The model-based system can fail due to its sensitivity to cognitive resources but it is more efficient in encoding previous experience of the agent. On the other hand, the model-free component is more affected by limited exploration but it is more reliable in its selections unaffected by the availability of cognitive resources. Consistent with the neural model, differentiations in behaviors among endophenotypes emerge in an inverted-U shape, where unbalanced control system are the most vulnerable to developing addiction.

The phenomenon of relapse is more elusive and the two models do not fully converge on this aspect. In the neural model, balanced dorsal and ventral endophenotypes respond well to both types of simulated treatments. For the unbalanced endophenotypes, however, only the appropriate treatment, targeting the dominant neural circuit, is effective. The simulations in the RL model do not show this same symmetric effects for the two treatments: the model-free treatment is effective for most endophenotypes, whereas the model-based treatment is mostly unsuccessful, with short relapse times across all endophenotypes, but the purely model-based one. The latter result is possibly due to the model learning process characterizing the model-based component, which is affected by conflicting information as drug use is associated with both positive and negative outcomes, experienced by the agent when entering the drug state under different conditions.

It is worth noting that habitual and goal-oriented behaviors have neural representations in the dorsal and ventral cortico-striatal circuits respectively, but they do not fully overlap with model-based and model-free control modalities in RL (Dolan and Dayan, 2013). Nonetheless, the neural and RL models independently simulate choices among competing options in addiction. Thus, we were able to test our hypothesis of endophenotypic differentiation under two complementary levels in Marr’s tri-level of analysis: the neural implementation and the algorithmic level (Marr and Poggio, 1976). This multilevel modeling approach has been often used in computational psychiatry (Maia and Frank, 2011; Montague et al., 2012; Adams et al., 2016; Hauser et al., 2016; Huys et al., 2016) to highlight model convergence and associate specific neural structure and dynamics with mathematical formalizations of optimal and suboptimal behavior in RL. The convergence of neural and RL model also provides more confidence in the reliability of the identified computational mechanisms underlying addiction and the associated characterization of endophenotypes. Importantly, the simulated endophenotype-dependent treatment effects can potentially inform the development of individualized treatments for addiction. For instance, both models indicate individuals showing unbalanced cortico-striatal activity or control modality are at higher risk of relapse and might need prolonged or ad hoc developed treatments, in comparison with balanced endophenotypes. Further studies will be required to provide empirical validation of our models. For example, connectivity analysis of fMRI imaging can be used to test effective connectivity among cortico-striatal circuits, in conjunction with cognitive tasks targeting the model-based and model-free control systems.

## Acknowledgments

The authors wish to thank Prof. Karl Friston for his comments and kind suggestions in shaping this manuscript.

## References

Adams RA, Huys QJ, Roiser JP (2016) Computational Psychiatry: towards a mathematically informed understanding of mental illness. J Neurol Neurosurg Psychiatry 87:53–63.

Afraimovich V, Tristan I, Huerta R, Rabinovich MI (2008) Winnerless competition principle and prediction of the transient dynamics in a Lotka-Volterra model. Chaos 18:043103.

Amit DJ (1989) Modeling brain function: the world of attractor neural networks. Cambridge; New York: Cambridge University Press.

Baldassarre G, Mannella F, Fiore VG, Redgrave P, Gurney K, Mirolli M (2013) Intrinsically motivated action-outcome learning and goal-based action recall: a system-level bio-constrained computational model. Neural Netw 41:168–187.

Balleine BW (2005) Neural bases of food-seeking: affect, arousal and reward in corticostriatolimbic circuits. Physiol Behav 86:717–730.

Balleine BW, O’Doherty JP (2010) Human and rodent homologies in action control: corticostriatal determinants of goal-directed and habitual action. Neuropsychopharmacology 35:48–69.

Belcher AM, Volkow ND, Moeller FG, Ferre S (2014) Personality traits and vulnerability or resilience to substance use disorders. Trends Cogn Sci 18:211–217.

Belin D, Everitt BJ (2008) Cocaine seeking habits depend upon dopamine-dependent serial connectivity linking the ventral with the dorsal striatum. Neuron 57:432–441.

Belin D, Deroche-Gamonet V (2012) Responses to novelty and vulnerability to cocaine addiction: contribution of a multi-symptomatic animal model. Cold Spring Harbor perspectives in medicine 2.

Belin D, Belin-Rauscent A, Everitt BJ, Dalley JW (2016) In search of predictive endophenotypes in addiction: insights from preclinical research. Genes, brain, and behavior 15:74–88.

Belin D, Mar AC, Dalley JW, Robbins TW, Everitt BJ (2008) High impulsivity predicts the switch to compulsive cocaine-taking. Science 320:1352–1355.

Belin D, Berson N, Balado E, Piazza PV, Deroche-Gamonet V (2011) High-novelty-preference rats are predisposed to compulsive cocaine self-administration. Neuropsychopharmacology 36:569–579.

Bjorklund A, Dunnett SB (2007) Dopamine neuron systems in the brain: an update. Trends Neurosci 30:194–202.

Carroll ME, Morgan AD, Lynch WJ, Campbell UC, Dess NK (2002) Intravenous cocaine and heroin self-administration in rats selectively bred for differential saccharin intake: phenotype and sex differences. Psychopharmacology (Berl) 161:304–313.

Cohen MX, Frank MJ (2009) Neurocomputational models of basal ganglia function in learning, memory and choice. Behavioural brain research 199:141–156.

Covington HE, 3rd, Miczek KA (2005) Intense cocaine self-administration after episodic social defeat stress, but not after aggressive behavior: dissociation from corticosterone activation. Psychopharmacology (Berl) 183:331–340.

Dalley JW, Everitt BJ, Robbins TW (2011) Impulsivity, compulsivity, and top-down cognitive control. Neuron 69:680–694.

Dalley JW, Fryer TD, Brichard L, Robinson ES, Theobald DE, Laane K, Pena Y, Murphy ER, Shah Y, Probst K, Abakumova I, Aigbirhio FI, Richards HK, Hong Y, Baron JC, Everitt BJ, Robbins TW (2007) Nucleus accumbens D2/3 receptors predict trait impulsivity and cocaine reinforcement. Science 315:1267–1270.

Daw ND, Dayan P (2014) The algorithmic anatomy of model-based evaluation. Philos Trans R Soc Lond B Biol Sci 369.

Daw ND, Niv Y, Dayan P (2005) Uncertainty-based competition between prefrontal and dorsolateral striatal systems for behavioral control. Nat Neurosci 8:1704–1711.

Daw ND, Gershman SJ, Seymour B, Dayan P, Dolan RJ (2011) Model-based influences on humans’ choices and striatal prediction errors. Neuron 69:1204–1215.

Dayan P (2009) Dopamine, reinforcement learning, and addiction. Pharmacopsychiatry 42 Suppl 1:S56–65.

Deco G, Jirsa VK, Robinson PA, Breakspear M, Friston K (2008) The dynamic brain: from spiking neurons to neural masses and cortical fields. PLoS Comput Biol 4:el000092.

desJardins ME, Durfee EH, Ortiz J, Charles L., Wolverton MJ (1999) A survey of research in distributed, continual planning. AI Magazine 20:13–22.

Dilleen R, Pelloux Y, Mar AC, Molander A, Robbins TW, Everitt BJ, Dalley JW, Belin D (2012) High anxiety is a predisposing endophenotype for loss of control over cocaine, but not heroin, self-administration in rats. Psychopharmacology (Berl) 222:89–97.

Dolan RJ, Dayan P (2013) Goals and habits in the brain. Neuron 80:312–325.

Doll BB, Daw ND (2016) The expanding role of dopamine. Elife 5.

Doll BB, Jacobs WJ, Sanfey AG, Frank MJ (2009) Instructional control of reinforcement learning: a behavioral and neurocomputational investigation. Brain Res 1299:74–94.

Draganski B, Kherif F, Kloppel S, Cook PA, Alexander DC, Parker GJ, Deichmann R, Ashburner J, Frackowiak RS (2008) Evidence for segregated and integrative connectivity patterns in the human Basal Ganglia. J Neurosci 28:7143–7152.

Economidou D, Pelloux Y, Robbins TW, Dalley JW, Everitt BJ (2009) High impulsivity predicts relapse to cocaine-seeking after punishment-induced abstinence. Biol Psychiatry 65:851–856.

Ersche KD, Turton AJ, Pradhan S, Bullmore ET, Robbins TW (2010) Drug addiction endophenotypes: impulsive versus sensation-seeking personality traits. Biol Psychiatry 68:770–773.

Everitt BJ, Robbins TW (2013) From the ventral to the dorsal striatum: devolving views of their roles in drug addiction. Neurosci Biobehav Rev 37:1946–1954.

Everitt BJ, Robbins TW (2016) Drug Addiction: Updating Actions to Habits to Compulsions Ten Years On. Annu Rev Psychol 67:23–50.

Fiore VG, Dolan RJ, Strausfeld NJ, Hirth F (2015) Evolutionarily conserved mechanisms for the selection and maintenance of behavioural activity. Philos Trans R Soc Lond B Biol Sci 370.

Fiore VG, Rigoli F, Stenner MP, Zaehle T, Hirth F, Heinze HJ, Dolan RJ (2016) Changing pattern in the basal ganglia: motor switching under reduced dopaminergic drive. Sci Rep 6:23327.

Fiore VG, Sperati V, Mannella F, Mirolli M, Gurney K, Friston K, Dolan RJ, Baldassarre G (2014) Keep focussing: striatal dopamine multiple functions resolved in a single mechanism tested in a simulated humanoid robot. Front Psychol 5:124.

Flagel SB, Waselus M, Clinton SM, Watson SJ, Akil H (2014) Antecedents and consequences of drug abuse in rats selectively bred for high and low response to novelty. Neuropharmacology 76 Pt B:425–436.

Flagel SB, Robinson TE, Clark JJ, Clinton SM, Watson SJ, Seeman P, Phillips PE, Akil H (2010) An animal model of genetic vulnerability to behavioral disinhibition and responsiveness to reward-related cues: implications for addiction. Neuropsychopharmacology 35:388–400.

Gannon BM, Galindo KI, Rice KC, Collins GT (2017) Individual Differences in the Relative Reinforcing Effects of 3,4-Methylenedioxypyrovalerone under Fixed and Progressive Ratio Schedules of Reinforcement in Rats. The Journal of pharmacology and experimental therapeutics 361:181–189.

Garrison KA, Potenza MN (2014) Neuroimaging and biomarkers in addiction treatment. Current psychiatry reports 16:513.

Gerfen CR, Surmeier DJ (2011) Modulation of striatal projection systems by dopamine. Annu Rev Neurosci 34:441–466.

Gershman SJ, Horvitz EJ, Tenenbaum JB (2015) Computational rationality: A converging paradigm for intelligence in brains, minds, and machines. Science 349:273–278.

Gillan CM, Kosinski M, Whelan R, Phelps EA, Daw ND (2016) Characterizing a psychiatric symptom dimension related to deficits in goal-directed control. Elife 5.

Gould RW, Duke AN, Nader MA (2014) PET studies in nonhuman primate models of cocaine abuse: translational research related to vulnerability and neuroadaptations. Neuropharmacology 84:138–151.

Grahn JA, Parkinson JA, Owen AM (2009) The role of the basal ganglia in learning and memory: neuropsychological studies. Behavioural brain research 199:53–60.

Gruber AJ, McDonald RJ (2012) Context, emotion, and the strategic pursuit of goals: interactions among multiple brain systems controlling motivated behavior. Front Behav Neurosci 6:50.

Gutkin BS, Dehaene S, Changeux JP (2006) A neurocomputational hypothesis for nicotine addiction. Proc Natl Acad Sci U S A 103:1106–1111.

Haber S (2008) Parallel and integrative processing through the Basal Ganglia reward circuit: lessons from addiction. Biol Psychiatry 64:173–174.

Haber SN (2003) The primate basal ganglia: parallel and integrative networks. J Chem Neuroanat 26:317–330.

Hauser TU, Fiore VG, Moutoussis M, Dolan RJ (2016) Computational Psychiatry of ADHD: Neural Gain Impairments across Marrian Levels of Analysis. Trends Neurosci 39:6373.

Hoffman RE, McGlashan TH (2001) Neural network models of schizophrenia. Neuroscientist 7:441–454.

Huys QJ, Maia TV, Frank MJ (2016) Computational psychiatry as a bridge from neuroscience to clinical applications. Nat Neurosci 19:404–413.

Hyman SE, Malenka RC, Nestler EJ (2006) Neural mechanisms of addiction: the role of reward-related learning and memory. Annu Rev Neurosci 29:565–598.

Jahanshahi M, Obeso I, Rothwell JC, Obeso JA (2015) A fronto-striato-subthalamic-pallidal network for goal-directed and habitual inhibition. Nat Rev Neurosci 16:719–732.

Jimenez VA, Grant KA (2017) Studies using macaque monkeys to address excessive alcohol drinking and stress interactions. Neuropharmacology 122:127–135.

Jonkman S, Pelloux Y, Everitt BJ (2012) Drug intake is sufficient, but conditioning is not necessary for the emergence of compulsive cocaine seeking after extended self-administration. Neuropsychopharmacology 37:1612–1619.

Jupp B, Dalley JW (2014) Behavioral endophenotypes of drug addiction: Etiological insights from neuroimaging studies. Neuropharmacology 76 Pt B:487–497.

Keramati M, Dezfouli A, Piray P (2011) Speed/accuracy trade-off between the habitual and the goal-directed processes. PLoS Comput Biol 7:e1002055.

Koob GF, Volkow ND (2016) Neurobiology of addiction: a neurocircuitry analysis. The lancet Psychiatry 3:760–773.

Maia TV, Frank MJ (2011) From reinforcement learning models to psychiatric and neurological disorders. Nat Neurosci 14:154–162.

Marr D, Poggio T (1976) From understanding computation to understanding neural circuitry. Cambridge: Massachusetts Institute of Technology, Artificial Intelligence Laboratory.

McClintick MN, Grant KA (2016) Aggressive temperament predicts ethanol self-administration in late adolescent male and female rhesus macaques. Psychopharmacology (Berl) 233:3965–3976.

Molander AC, Mar A, Norbury A, Steventon S, Moreno M, Caprioli D, Theobald DE, Belin D, Everitt BJ, Robbins TW, Dalley JW (2011) High impulsivity predicting vulnerability to cocaine addiction in rats: some relationship with novelty preference but not novelty reactivity, anxiety or stress. Psychopharmacology (Berl) 215:721–731.

Montague PR, Dolan RJ, Friston KJ, Dayan P (2012) Computational psychiatry. Trends Cogn Sci 16:72–80.

Moore A, Atkeson CG (1993) Prioritized sweeping: Reinforcement learning with less data and less time. Machine Learning 13:103–130.

Morgan D, Grant KA, Gage HD, Mach RH, Kaplan JR, Prioleau O, Nader SH, Buchheimer N, Ehrenkaufer RL, Nader MA (2002) Social dominance in monkeys: dopamine D2 receptors and cocaine self-administration. Nat Neurosci 5:169–174.

Nadal R, Armario A, Janak PH (2002) Positive relationship between activity in a novel environment and operant ethanol self-administration in rats. Psychopharmacology (Berl) 162:333–338.

Nader MA, Czoty PW (2005) PET imaging of dopamine D2 receptors in monkey models of cocaine abuse: genetic predisposition versus environmental modulation. Am J Psychiatry 162:1473–1482.

Nestler EJ, Aghajanian GK (1997) Molecular and cellular basis of addiction. Science 278:58-63.

Obeso JA, Rodriguez-Oroz MC, Stamelou M, Bhatia KP, Burn DJ (2014) The expanding universe of disorders of the basal ganglia. Lancet 384:523–531.

Pelloux Y, Everitt BJ, Dickinson A (2007) Compulsive drug seeking by rats under punishment: effects of drug taking history. Psychopharmacology (Berl) 194:127–137.

Pelloux Y, Murray JE, Everitt BJ (2015) Differential vulnerability to the punishment of cocaine related behaviours: effects of locus of punishment, cocaine taking history and alternative reinforcer availability. Psychopharmacology (Berl) 232:125–134.

Perry JL, Carroll ME (2008) The role of impulsive behavior in drug abuse. Psychopharmacology (Berl) 200:1–26.

Pezzulo G, Rigoli F, Chersi F (2013) The mixed instrumental controller: using value of information to combine habitual choice and mental simulation. Front Psychol 4:92.

Piazza PV, Deminiere JM, Le Moal M, Simon H (1989) Factors that predict individual vulnerability to amphetamine self-administration. Science 245:1511–1513.

Piray P, Keramati MM, Dezfouli A, Lucas C, Mokri A (2010) Individual differences in nucleus accumbens dopamine receptors predict development of addiction-like behavior: a computational approach. Neural Comput 22:2334–2368.

Rabinovich MI, Varona P, Selverston AI, Abarbanel HDI (2006) Dynamical principles in neuroscience. Reviews of Modern Physics 78.

Redish AD (2004) Addiction as a computational process gone awry. Science 306:1944–1947.

Redish AD, Jensen S, Johnson A (2008) A unified framework for addiction: vulnerabilities in the decision process. Behav Brain Sci 31:415–437; discussion 437-487.

Sadacca BF, Jones JL, Schoenbaum G (2016) Midbrain dopamine neurons compute inferred and cached value prediction errors in a common framework. Elife 5.

Schultz W, Dayan P, Montague PR (1997) A neural substrate of prediction and reward. Science 275:1593–1599.

Silver D, Huang A, Maddison CJ, Guez A, Sifre L, van den Driessche G, Schrittwieser J, Antonoglou I, Panneershelvam V, Lanctot M, Dieleman S, Grewe D, Nham J, Kalchbrenner N, Sutskever I, Lillicrap T, Leach M, Kavukcuoglu K, Graepel T, Hassabis D (2016) Mastering the game of Go with deep neural networks and tree search. Nature 529:484–489.

Simon DA, Daw ND (2012) Dual-system learning models and drugs of abuse. In: Computational Neuroscience of Drug Addiction, 1 Edition (Gutkin B, Ahmed SH, eds), pp 145–161. New York: Springer-Verlag.

Singh S, Jaakkola T, Littman ML, Szepesvári C (2000) Convergence Results for Single-Step On-Policy Reinforcement-Learning Algorithms. Machine Learning 38:287–308.

Smethells JR, Zlebnik NE, Miller DK, Will MJ, Booth F, Carroll ME (2016) Cocaine self-administration and reinstatement in female rats selectively bred for high and low voluntary running. Drug and alcohol dependence 167:163–168.

Suto N, Austin JD, Vezina P (2001) Locomotor response to novelty predicts a rat’s propensity to self-administer nicotine. Psychopharmacology (Berl) 158:175–180.

Sutton RS (1990) Integrated architecture for learning, planning, and reacting based on approximating dynamic programming. In: Proceedings of the seventh international conference (1990) on Machine learning, pp 216–224. Austin, Texas, USA: Morgan Kaufmann Publishers Inc.

Sutton RS, Barto AG (1998) Reinforcement Learning: An Introduction. Cambridge, MA: MIT Press.

Verdejo-Garcia A, Lawrence AJ, Clark L (2008) Impulsivity as a vulnerability marker for substance-use disorders: review of findings from high-risk research, problem gamblers and genetic association studies. Neurosci Biobehav Rev 32:777–810.

Voikow ND, Morales M (2015) The Brain on Drugs: From Reward to Addiction. Cell 162:712–725.

Volkow ND, Fowler JS, Wang GJ, Swanson JM, Telang F (2007) Dopamine in drug abuse and addiction: results of imaging studies and treatment implications. Archives of neurology 64:1575–1579.

Voon V, Reiter A, Sebold M, Groman S (2017) Model-Based Control in Dimensional Psychiatry. Biol Psychiatry 82:391–400.

Watkins CJ, Dayan P (1992) Q-Learning. Machine Learning 8:279–292.

Willuhn I, Burgeno LM, Everitt BJ, Phillips PE (2012) Hierarchical recruitment of phasic dopamine signaling in the striatum during the progression of cocaine use. Proc Natl Acad Sci U S A 109:20703–20708.

Yin HH, Knowlton BJ, Balleine BW (2004) Lesions of dorsolateral striatum preserve outcome expectancy but disrupt habit formation in instrumental learning. Eur J Neurosci 19:181–189.

